# Fission or fusion: shoaling adaptations in green chromides (*Etroplus suratensis*) across multiple manipulations

**DOI:** 10.1101/2025.06.24.661235

**Authors:** Chena Desai, Dinesh Nariani, Rudrik Dave, Ratna Ghosal

**Affiliations:** Biological and Life Sciences, School of Arts and Sciences, Ahmedabad University, Ahmedabad, Gujarat, India; Mathematical and Physical Sciences, School of Arts and Sciences, Ahmedabad University, Ahmedabad, Gujarat, India; Department of Zoology, School of Sciences, Gujarat University, Ahmedabad, Gujarat, India

**Keywords:** Behavioural plasticity, Collective behaviour, fission-fusion, green chromides, resource constraint, shoal

## Abstract

Collective behaviour contributes towards increased fitness, however, in fission-fusion societies, the decision to participate in a group is based on cost-benefit ratio derived under a given condition. In today’s world, this ratio dramatically changes due to increased challenges in degraded habitats, impacting behavioural decisions towards social grouping. In this paper, we used fish shoal as a unit of collective behaviour, and investigated variations in shoaling adaptations across a range of manipulations, mimicking challenges faced by the species under natural conditions. We used green chromides (*Etroplus suratensis*), a cichlid fish species, and characterised their shoaling behaviour under laboratory conditions for two group sizes, 4 and 8. We then examined the effects of different manipulations, for example, starvation and reduced space (both mimicking resource constrained conditions), and presence of hetero species including tilapia (*Oreochromis* sp., an alien species widespread within the habitats of *E. suratensis*) on their shoaling behaviour. Our results showed that *E. suratensis* formed shoals in both group sizes, 4 and 8. In subsequent analyses, we used group size 8 as a control and demonstrated that space reduction and starvation significantly impacted shoaling, reducing shoal splits and occurrences of solitary fish, while increasing incidences of single, cohesive shoals comprising all individuals. *E. suratensis* also formed mixed-species shoals with alien *Oreochromis* spp., but exhibited a large percentage (in absolute terms) of shoal splits. Overall, *E. suratensis* exhibited plasticity in their shoaling behaviour, and increased fusion of shoals under challenging conditions, which was in contrast to showing more fission in presence of the alien hetero species.

## 1. Introduction

Group formation or collective behaviour is a widely prevalent strategy across diverse taxonomic groups, and provides several fitness benefits including predator defence, improved food acquisition, collective vigilance, social learning, mating opportunities, and optimizing energy expenditure during locomotion (Ioannou, et al. 2011; Johannesen et al. 2014; Magurran, 1990; Parrish & Edelstein-Keshet, 1999; Pitcher & Parrish, 1993; Ward et al. 2011). However, in most fission-fusion societies, grouping behaviour is highly dynamic in nature with organisms exhibiting informed decision-making based on their perception of the immediate local environment (Aguilar-Melo et al. 2018; Aureli et al. 2008; Kerth et al. 2006). Such dynamicity is highly adaptive in nature, particularly under the current circumstances, wherein the natural environment of a species is highly threatened by increased trends of urbanization, leading to pollution, deteriorating the quality of the habitats and increasing the occurrences of non-native (alien) species (Simkin et al. 2022; Larrañaga et al. 2019; MacGregor and Ioannou 2023). Studies have shown that when faced with severe challenges, organisms may either decrease (increased fusion) the fluidity in social groupings, possibly seeking cooperation to combat adverse conditions, or may split (fission) the groups, potentially exhibiting social conflict among the group members (Tiddy et al. 2024; Aureli et al. 2008). Such flexibility showcases plasticity of the behavioural trait and demonstrates the various ways in which organisms strike a fine balance between the cost and the benefits derived under the collective behavioural regime (Ginnaw et al. 2020; Li et al. 2022; Tiddy et al. 2024).

With regards to fish taxa, group behaviour and collective motion has been studied in great detail, documenting these ectotherms to be extremely sensitive, exhibiting dynamic strategies in their collective motion in order to adapt to their immediate local environment (Cooper et al. 2018; Killen et al. 2016). Group behaviour in fish has been defined in two ways, shoaling and schooling. Shoaling represents a more social structure and schooling is more of a coordinated physical movement (Pitcher and Parrish, 1993). A shoal can also be defined as a multilayered structure, similar to a multilayer perceptron (Popescu et al. 2009). Each layer of the shoal contributes towards different aspects of group dynamics, wherein the outer layers, for example, inter-individual distance, individual speed, shoal size, polarization, and nearest-neighbour distance, represent observable components (referred to as structural components) that progressively shape the overall structure of the shoal. Beneath these outer layers are the hidden proximate layers that integrate sensory inputs, cognitive abilities, and physiological state of individuals in order to contribute towards decision-making processes during shoal formation. Fish adjust different components of the outer layers in response to various environmental conditions (for example, predator presence and resource availability) via continuous feedback from the proximate layers. This cross-talk between outer and inner layers enables fish shoals to adapt to changing environmental and ecological conditions, resulting in dynamic variations across time and space.

The rate of change taking place in the social environment of a species has drastically increased over the years, mainly because of the changes in their physical environment due to urbanization, reducing the quality of the habitats, and globalization that has led to increased competition due to high rate of introductions of non-native (alien or exotic) species (Bernery et al. 2024; Lawson et al. 2024; Xu et al. 2024). To understand how such changes impact the fission-fusion dynamics in fish shoals, studies have focussed on investigating shoaling adaptations under a range of different ecological and environmental conditions. Such adaptations showcase cognitive skills of the organisms towards perceiving challenges in their local environment, and in turn, making informed decisions to maximize fitness under the given condition (Michael et al. 2021; Herbert et al. 2017; Tiddy et al. 2024). Most studies showed that in order to adapt, species modify structural components (the outer layer) of shoals, for example, group size, inter-individual distance (IID), nearest neighbour distance (NND), and shoal speed, and such modifications may vary across a range of ecological and environmental conditions (Krause et al. 2000; Ginnaw et al. 2020; Bhat et al. 2015). For example, a study by MacGregor and Ioannou (2023) on three-spined sticklebacks *Gasterosteus aculeatus* Linnaeus 1758 found that shoals reduced their nearest neighbour distance (NND) and swimming speed in turbid conditions when compared to clear waters. Turbidity can impair sensory modalities, making effective information transfer within shoals more challenging. The authors speculated that higher shoal cohesion led to increased fusion, enhancing transfer of information, while decreasing speed might have helped to maintain cohesiveness in the shoals (Chamberlain and Ioannou, 2019; Kimbell and Morrell, 2015). Similarly, resource constraints, for example, reduced availability of dissolved oxygen in eutrophic habitats and limitations in finding food under conditions of high competition (that may even lead to starvation) have also been shown to influence shoaling behaviours of fishes (Domenici et al. 2017; Krause, 1993; Hansen et al. 2015; Wilson et al. 2019; Zheng and Fu, 2021). For example, Iberian barbel *Luciobarbus bocagei* Steindachner 1864 formed less cohesive shoals, thus increasing fission under oxygen depleted conditions when compared to the control group (shoals in well-oxygenated water), possibly to minimize crowding and competition for oxygen, and in turn, reducing energy expenditure under such unfavourable conditions (Hayes et al. 2019).

In addition to the changes in the abiotic environment, introduction of alien species may also have a potential impact on the shoaling behaviours of the native species, wherein the alien and the native species could be behaviourally novel to each other due to non-overlapping natural distributions (Ali et al. 2018). Additionally, alien species may often derive fitness advantages by shoaling with the natives, as the latter are far more familiar within the given environment (Camacho-Cervantes et al. 2014a). For example, Camacho-Cervantes et al. (2014b) investigated shoaling behaviour of guppies *Poecilia reticulata* Peters 1859, one of the world’s most invasive freshwater fish that has successfully invaded several freshwater ecosystems in Mexico) along with four different species of topminnows-*Skiffia bilineata* Bean 1887, *Zoogoneticus tequila* Webb and Miller 1998, *Xenotoca eiseni* (Rutter 1896) and *Girardinichthys viviparous* Bustamante 1839 that are native to Mexico. The study showed that guppies shoaled with *S. bilineata, Z. tequila, X. eiseni*, with no significant differences in NND between conspecific (conspecific guppy shoals) and mixed-species shoals, and additionally, the presence of native topminnows increased the foraging efficiency of the guppies. Contrastingly, *P. reticulata* did not shoal (with significant differences in NND when compared to conspecific *P. reticulata* shoals) with the topminnow species, *G. viviparous,* and interestingly, its presence did not enhance the foraging efficiency of *P. reticulata* (Camacho-Cervantes et al. 2014b). Thus, based on cost-benefit ratio across different environments, fishes modify their shoaling behaviour, via adjustments of their structural components, and such plasticity helps maintain the dynamics of collective behaviour, contributing towards the overall fitness of the shoaling individual (Ioannou and Laskowski, 2023).

In this paper, we aimed to examine shoaling behaviour of green chromides *Etroplus suratensis* (Bloch 1790), a freshwater fish species belonging to the Cichlidae family, and are endemic to India and the subcontinent (Alex et al. 2020; Mallick et al. 2022; Desai and Ghosal, 2024). *E. suratensis* are members of the subfamily Etroplinae, which represents one of the least-studied groups of ancient cichlids (Matschiner, 2019). *E. suratensis* show a range of complex behaviours, for example, courtship and parental care, and inhabit different environmental conditions, including lentic and lotic ecosystems, and in both turbid and clear water conditions, along the east and west coasts of India (Alex et al. 2020; Padmakumar et al. 2012). Most of the native habitats of the *E. suratensis* face severe threats, for example, urbanization due to port construction, environmental degradation from tourism, and intense resource competition due to introduction of several non-native species, for example, tilapia *Oreochromis* spp., three-spotted gourami *Trichogaster trichopterus* (Pallas 1770), suckermouth catfish *Pterygoplichthys* spp., African sharptooth catfish *Clarias gariepinus* (Burchell 1822), common carp *Cyprinus carpio* Linnaeus 1758, and mosquitofish *Gambusia affinis* Baird & Girard 1853 (Raj et al. 2021; Roshni et al. 2021). We hypothesised that challenges faced by the *E. suratensis* in their local environment may impact their behavioural interactions, in particular, their shoaling dynamics. With this background, we aimed to study shoaling behaviour of the *E. suratensis* under laboratory conditions and subsequently investigated their behavioural adaptations under a range of different environmental conditions, mimicking challenges that the species face in their natural environment, for example, reduced space availability due to habitat fragmentation, physiologically starved conditions due to lower abundance of food in deteriorated habitats, and presence of alien hetero species due to large-scale introduction of exotic fishes in freshwater habitats inhabited by the *E. suratensis*. Overall, the study will contribute towards understanding shoaling dynamics of *E. suratensis*, while providing insights into the adaptations of collective behaviour in an early-diverging cichlid.

## 2. Methods

### 2.1. Ethical statement

All the necessary ethical approvals for this study were obtained from the University Research Board and the Institute Biosafety Committee under the project code AU/SUG/SAS/BLS/2018-19/20.

### 2.2. Study animals

*E. suratensis* and orange chromide *Pseudetroplus maculatus* (Bloch 1795) were collected using a cast net from the Hooghly River, near Mousuni Island in West Bengal, India (21°41’44.88’’ N, 88°11’8.714’’ E), during July and August 2022. *Oreochromis* spp. were obtained from the Sabarmati River in Ahmedabad, Gujarat, India (22°59’51.3492’’N, 72°33’51.6132’’ E), in December 2022. All three species were transported to the fish laboratory at Ahmedabad University in India, for subsequent experiments. Transport of the fishes was done via airfreight in temperature-controlled containers, and with optimum levels of dissolved oxygen. The fish were housed in 70-liter glass tanks and fed commercially available Growfin feed (Growel, Andhra Pradesh, India) at a rate of 5% of their body weight, once daily. The water temperature in the tanks was maintained at 27±2°C, having a day-night cycle of 14:10 hours. Each species was housed separately and acclimatized for at least six months before the start of the experiments. Fish were screened visually for parasites and fin rot, and only healthy fish were selected for experiments. *E. suratensis* (green chromide, referred to as GC from here onwards), *P. maculatus* (orange chromide, referred to as OC from here onwards), and *Oreochromis* spp. (tilapia, referred to as T from here onwards) had a total length [Mean±standard error (SE)] [from the tip of the snout to the tip of the caudal fin (Önsoy et al. 2011)] of 5.5 ± 0.5 cm. *Oreochromis* spp. of a larger body size (total length = 10±0.7 cm, abbreviated as LT from here onwards) were also used in the experiments. No mortality of fish was recorded during the entire experimental duration.

### 2.3. Experimental setup and protocol for shoaling trials

A custom-built glass tank (dimensions: 120 cm × 120 cm × 22.5 cm) lined with polyvinyl chloride (PVC) sheet was used for all the experiments. The edges of the tank were sealed with PVC sheets to prevent fish from gathering in the corners. An overhead camera (Vivotek, Taiwan) was mounted 120 cm above the tank to record the trials. The tank was filled with water to a depth of 6 cm, this depth ensured that no two fish could overlap vertically, facilitating the measurements for shoaling components. The experiments were conducted under low light and low sound conditions to minimize external disturbances. Different treatments were applied based on the specific research questions (Table 1), and six biological replicates were performed for each trial type (Table 1). For the hetero species treatment, four GC were combined with four individuals of hetero species, including either OC (setup number 3), or T (setup number 4), or LT (setup number 5). For the starvation treatment, individuals were starved for either 24 or 48 hours before the experiment, depending on the trial type. In the 24-hour starvation setup (setup number 6), the tank was cleaned, and no food was provided for the next 24 hours. The same process was followed for the 48-hour starvation setup (setup number 7). In both cases, food was provided only after the experiments. To manipulate space availability (setup number 8), we reduced the available space to 50% by making a partition in the middle of the experiment tank using PVC sheets. In the reduced space setup, the ratio of area occupied by eight fish:total area available was approximately 1:25.3. In contrast, in all other setups, this ratio was 1:50.6, reflecting greater spatial availability under unmanipulated conditions. Each trial included a 30-minute acclimatization period followed by a 5-minute observation phase. All trials were conducted in the morning, between 0800 and 1100 am.

**Table 1.** Schematic representation of experimental setups.

| Setup number<sup>a</sup> | Treatments | Trial-type | Details |
| --- | --- | --- | --- |
| 1 | Con species | 4GC | 4 GC |
| 2 |  | 8GC | 8 GC |
| 3 | Hetero species | GC+OC | 4 GC + 4 OC |
| 4 |  | GC+T | 4 GC + 4 T |
| 5 |  | GC+LT | 4 GC + 4 LT |
| 6 | Starvation | GC_24H | 8 GC after 24-hour (24H) starvation |
| 7 |  | GC_48H | 8 GC after 48-hour (48H) starvation |
| 8 | Space reduction | GC_SR | 8 GC in reduced space (SR) |

<sup>a</sup>Setups 1 and 2 were used to address the first objective, while setups 2–8 were used to address the second objective.

### 2.4. Data processing

Videos were converted to MPEG-4 format using Microsoft Clipchamp, and the x and y coordinates of the fish were tracked using UMA Tracking software (Yamanaka and Takeuchi, 2018). All the tracking errors were manually corrected using the UMA Tracking Corrector software (Yamanaka and Takeuchi, 2018). Each 5-minute trial consisted of 9,000 frames, captured at a rate of 30 frames per second. Details on the methods of measuring each of the shoaling components, including IID, shoal size, solitary fish, and speed, are described below. To calculate the shoaling components (except speed), we analysed every 30th frame, resulting in 301 frames per trial. Due to logistical constraints, individual fish could not be identified throughout the video. However, species (GC, OC, T, and LT) level identification was possible for the experimental trials.

### 2.5. Readouts for different structural components

We used several readouts to quantify different structural components of a shoal, ranging from global parameters of collective behaviour, for example, inter-individual distance (IID), to local parameters of shoaling patterns, for example, shoal size, solitary fish, shoal splits and speed (Rieucau et al. 2015; Allibhai et al. 2023).

#### 2.5.1. Inter-individual distance (IID)

Inter-individual distance (IID) was calculated by measuring distances between a given fish pair, and then averaging across all pairs (Allibhai et al. 2023; Luo et al. 2024), including all the frames (N=301) in a trial, providing a single average value for a trial (Desai et al. 2025).

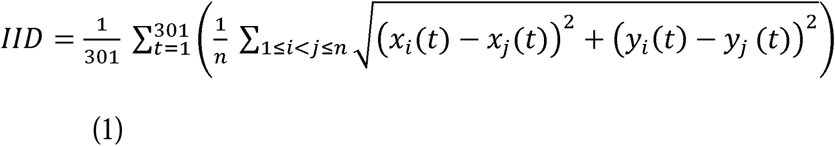

where x_i_ and y_i_ are the horizontal and vertical positions of fish i; x_j_ and y_j_ are the positions of fish j in the shoal at time t and n is the total number of unique fish pairs in a frame.

#### 2.5.2. Shoal size (number of individuals in a shoal)

Based on the nearest neighbour distance (NND) criterion, we classified individuals to be shoaling when they occurred within NND ≤4 body lengths (Pitcher and Parrish, 1993; Mallick et al. 2022), and accordingly calculated the shoal size. For comparative analyses across different treatments, we considered only the percentage of frames that had the maximum shoal size that is all individuals of a given treatment formed one shoal. For each trial, IID (as described above, distance between a given pair of individuals, Fu, 2016) was calculated for each frame (N=301) and then binarized based on the defined thresholds: distances ≤4 body lengths were marked as 1 (shoal) and >4 body lengths as 0. This process generated 301 CSV files per trial, each representing a single frame with binary data for each pair of individuals. The generated CSV files were then analysed using a Python script (Desai et al. 2025) to calculate the percentage of frames having the maximum shoal size.

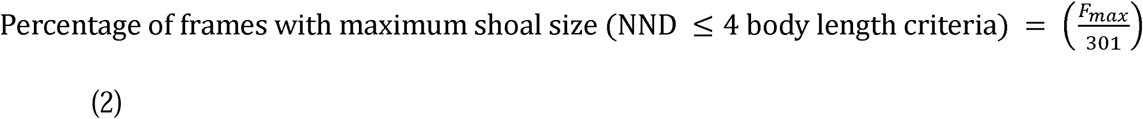

where F_max_ is the total number of frames in which all the fish formed a single shoal based on ≤4 body lengths criteria.

#### 2.5.3. Shoal cohesion

Shoal cohesion was considered when NND ≤1 body length (Pitcher and Parrish, 1993; Lucon-Xiccato et al. 2022). Maximum shoal size (all individuals of a given treatment formed one shoal) was calculated in a similar fashion as described above, for the NND ≤4 body length category.

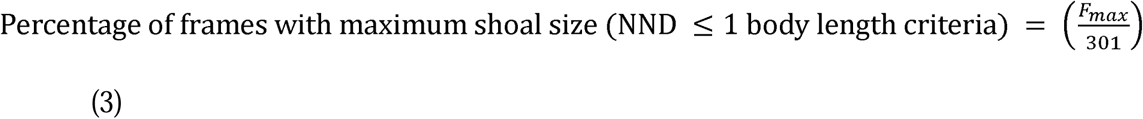

where F_max_ is the total number of frames in which all the fish formed a single shoal based on ≤1 body lengths criteria.

#### 2.5.4. Splits in the shoal

Shoal split was quantified as per Equation 1 for all the frames (N=301) for a given trial (Ghosal et al. 2016; Mallick et al. 2022):

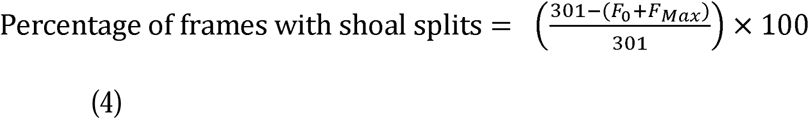

where F_0_ is the total number of frames with 0 shoals and F_max_ is the total number of frames in which all the fish formed a single shoal.

#### 2.5.5. Solitary fish

The number of frames with solitary individuals, fish that were not part of the shoal (as per the NND ≤4 body length criteria), was calculated for each trial and was reported as the percentage of frames containing at least one solitary fish for a trial for a given treatment (Ghosal et al. 2016).

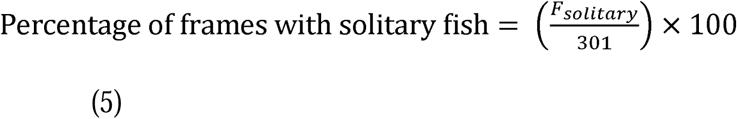

where F_solitary_ is the total number of frames containing at least one solitary fish.

#### 2.5.6. Speed

To calculate the speed of individual GC, we used the tracking software FishyGrid (Sommer-Trembo et al. 2020), which divided the entire bottom area of the tank into a grid, wherein each cell represented a square having a dimension of one body length (total length) of the fish. The entire dimension of the grid was 21 (X-axis) by 7 cells (Y-axis). For each trial of duration 5 minutes, we manually tracked three randomly selected GC for one minute at intervals of 0-1, 2-3, and 4-5 minutes of the trial, and during the tracking, the FishyGrid software recorded x-y coordinates that were occupied by the fish. The coordinates covered by the fish during the tracking were then converted into real-world distances while using the body length of the fish as the reference scale. Speed was then calculated as the distance travelled (in cm, units of body length) divided by the time (in seconds) taken (Tang et al. 2017; Luo et al. 2024).

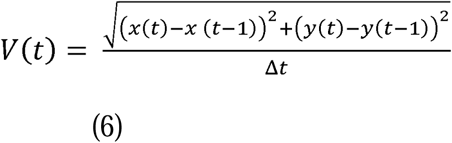

where V(t) is the swimming speed of an individual fish in cm/s; x(t) and y(t) are the fish’s horizontal and vertical positions in the grid at time t, and x(t−1) and y(t−1) are the positions at the previous time t−1; Δt is the time difference between these two points.

### 2.6. Statistical analysis

All readouts were checked for normality using the Shapiro-Wilk test. Data that met the normality assumption were analysed using parametric tests, while non-parametric methods were used for data that did not. Homogeneity of variance was assessed for all the readouts using the F-test. To examine if GC shoaled under laboratory conditions and whether the shoaling components varied across different shoal sizes or not, we did pairwise comparisons of each of the structural components between 4GC and 8GC trial types. To assess the effect of different treatments on the shoaling dynamics of GC, six separate One-way ANOVAs or Kruskal-Wallis tests (depending on normality tests) were conducted for each shoaling component: IID, maximum shoal size (as per NND≤4 body length criterion), shoal cohesion (as per NND ≤1 body length criterion), shoal splits, solitary fish, and speed, while comparing across treatments (setups 2 to 8; Table 1). When significant effects were detected, post-hoc Dunnett’s or Dunn’s tests with Bonferroni corrections were performed to compare each trial type with the 8GC trial type, which was considered as a control (with no manipulations). Data were analysed in R (version 4.3.2), with the significance level set at 0.05.

## 3. Results

Table 2 provides the details on mean±SE values for each of the structural components for all the setups (1-8).

**Table 2.**
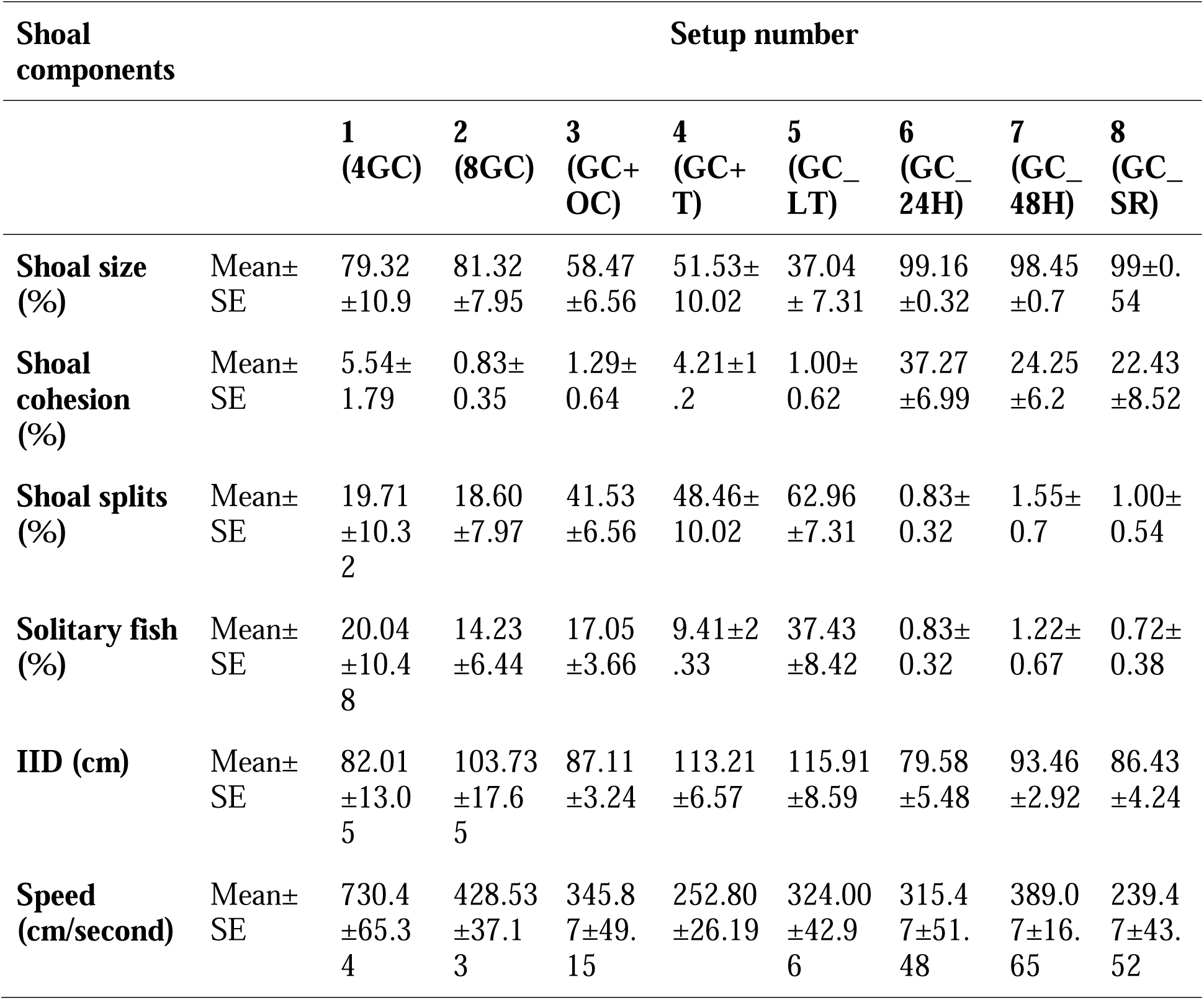
Mean ± SE for each shoal component across different trial types.

### 3.1. Conspecific shoaling

GC predominantly formed a single shoal including all individuals, for both the tested group sizes, 4GC and 8GC. Percentage of frames having the maximum shoal size (based on NND ≤4 body length criterion) (Wilcoxon rank sum exact test: *W* = 20, *P* = 0.8182; *N* = 12; Figure 1a), shoal splits (Student’s t-test: *t* = −0.26, *P* = 0.799; *N* = 12; Figure 1c), solitary fish (Student’s t-test: *t* = −0.692, *P* = 0.5045; *N* =12; Figure. 1d) and IID (Student’s t-test: *t* = −0.989, *P* = 0.346; *N* = 12; Figure 2a), did not differ significantly across the tested group sizes, 4GC and 8GC. However, speed and shoal cohesion were significantly different across the group sizes. 4GC exhibited a greater percentage of cohesive shoals (Student’s t-test, t = 2.73, P = 0.021; *N* = 12; Figure 1b) and significantly higher speed (Student’s t-test, *t* = 3.99, *P* = 0.0025; *N* = 12; Figure 2b) when compared to 8GC. 8GC was regarded as a control in further experiments, as it served as a testable group size for different manipulations, for example, the introductions of hetero species.

**Figure 1.**
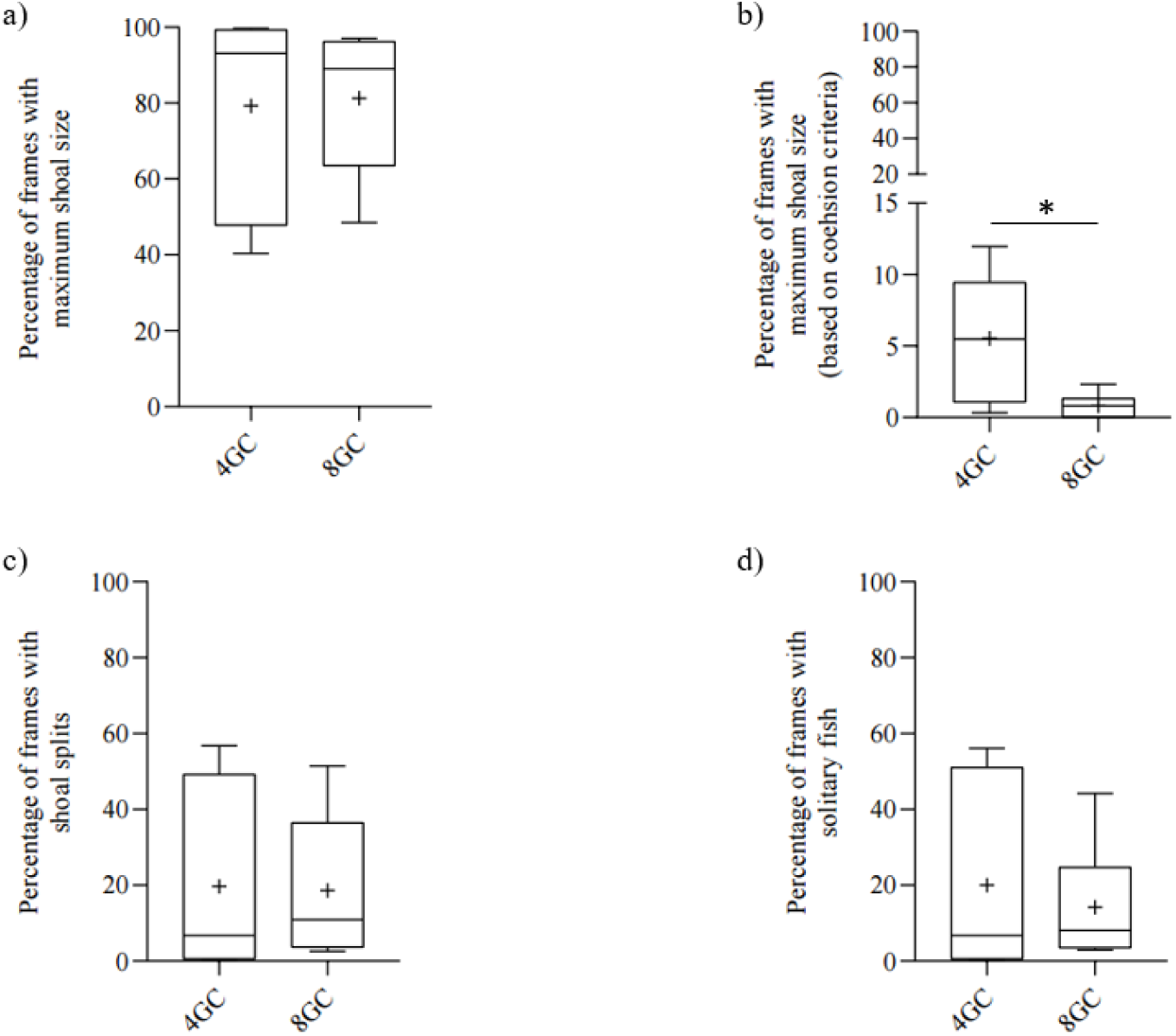
Effect of group size on (a) percentage of frames with maximum shoal size (based on NND ≤4 body length criteria), (b) percentage of frames with maximum shoal size (based on cohesion criteria: NND ≤1 body length criteria), (c) percentage of frames with shoal splits, and (d) percentage of frames with solitary fish, readouts for shoaling behaviour of GC (setup 1 and 2). The black center line inside each box represents the median (50th percentile), the box denotes the interquartile range (25th–75th percentiles), and the whiskers indicate the minimum and the maximum values. The ‘+’ symbol represents the mean values and * indicates significant (P<0.05) differences between 4GC and 8GC.

**Figure 2.**
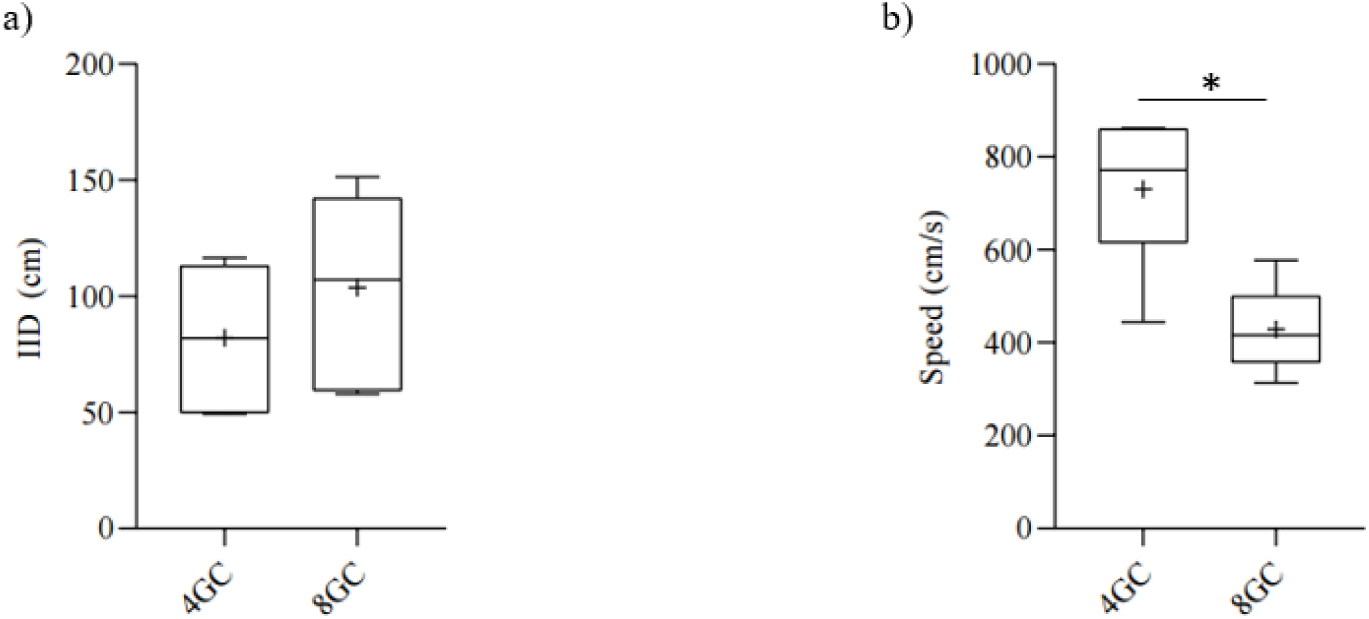
Effect of group size on (a) Inter-individual distance (IID), and (b) speed (cm/s), readouts for shoaling behaviour of GC (setup 1 and 2). The black centre line inside each box represents the median (50th percentile), the box denotes the interquartile range (25th–75th percentiles), and the whiskers indicate the minimum and the maximum values. The ‘+’ symbol represents the mean values and *indicates significant (P<0.05) differences between 4GC and 8GC.

### 3.2. Conspecific shoaling under different treatments

All the treatments, including starvation, space availability and presence of hetero species, had a significant effect (Figure 3 and 4) on multiple structural components (maximum shoal size: Kruskal Wallis: *H_6_* = 32.95, *P* < 0.0001, *N* = 42; shoal cohesion: Kruskal Wallis: *H_6_* = 33.26, *P* < 0.0001, *N* = 42; shoal split: Kruskal-Wallis: *H_6_* = 32.95, *P* < 0.0001, *N* = 42; solitary fish: Kruskal-Wallis: *H_6_* = 30.67, *P* < 0.0001, *N* = 42; speed: one-way ANOVA: *F_6,35_*= 2.912, *P* = 0.021, *N* = 42; IID: Kruskal Wallis: *H_6_* = 16.20, *P* = 0.0127, *N* = 42). Post hoc comparisons showed that GC_24H and GC_SR formed a significantly larger percentage of the maximum shoal size [as per NND ≤4 body length criteria (GC_24H-Dunn’s test: *P* = 0.03 and GC_SR-Dunn’s test: *P* = 0.0046); Figure 3a] and formed significantly more cohesive shoals [as per NND ≤1 body length criteria (GC_24H-Dunn’s test: *P* = 0.03 and GC_SR-Dunn’s test: *P* = 0.029) Figure 3b] when compared to 8GC. Post hoc tests also showed a significantly lower percentage of shoal splits (GC_24H-Dunn’s test: *P* = 0.03 and GC_SR-Dunn’s test: *P* = 0.047; Figure 3c) and solitary fish (GC_24H-Dunn’s test: *P* = 0.03 and GC_SR-Dunn’s test: *P* = 0.028; Figure 3d) in GC_SR and GC_24H, each when compared to 8GC. However, GC_48H had significant differences only in percentage of cohesive shoals (NND ≤1 body length criteria; Dunn’s test: *P* = 0.03) when compared to 8GC. No significant differences (post-hoc Dunnett’s test) were obtained for maximum shoal size, shoal split and solitary fish for GC_48H when compared to 8GC. Similarly, no significant differences (post-hoc Dunnett’s test) were obtained in speed for both GC_24H and GC_48H, each when compared to 8GC. Contrastingly, GC_SR exhibited significantly lower speed than 8GC (Dunnett’s test: *P* = 0.049; Figure 4b).

**Figure 3.**
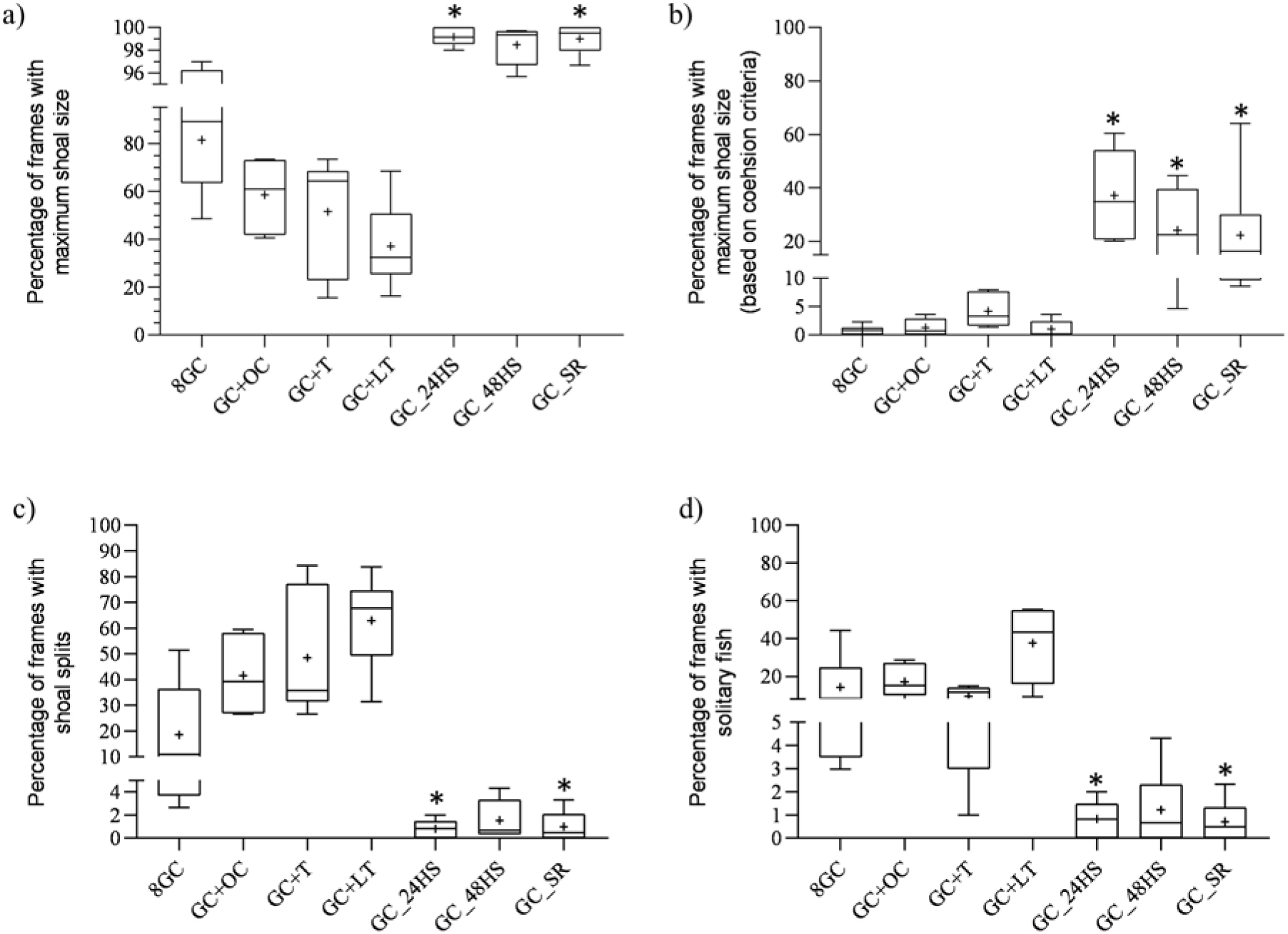
Effect of different treatments on (a) percentage of frames with maximum shoal size (as per NND ≤4 body length criteria), (b) percentage of frames with shoal cohesion (maximum shoal size as per NND ≤1 body length criteria), (c) percentage of frames with shoal splits, and (d) percentage of frames with solitary fish, readouts for shoaling behaviour of GC (setups 2 to 8). The black center line inside each box represents the median (50th percentile), the box denotes the interquartile range (25th–75th percentiles), and the whiskers indicate the minimum and the maximum values. The ‘+’ symbol represents the mean values and *indicates significant (P<0.05) differences between a given trial type and 8GC. Data for 8GC is repeated from Figure 1 for comparison purposes only.

**Figure 4.**
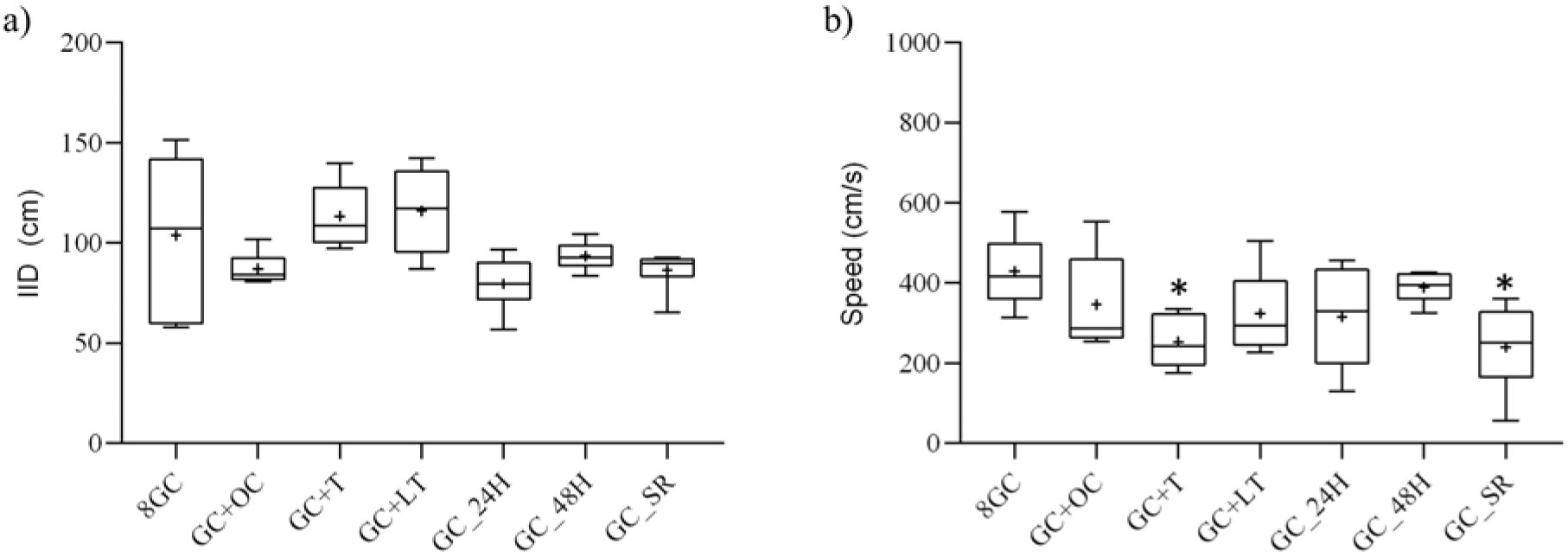
Effect of different treatments on (a) inter-individual distance (IID) (b) speed (cm/s), readouts for shoaling behaviour in GC (setups 2 to 8). The black center line inside each box represents the median (50th percentile), the box denotes the interquartile range (25th–75th percentiles), and the whiskers indicate the minimum and the maximum values. The ‘+’ symbol represents the mean values and *indicates significant (P<0.05) differences between a given trial type and 8GC. Data for 8GC is repeated from Figure 2 for comparison purposes only.

None of the structural components (maximum shoal size, cohesion, shoal split and solitary fish), except speed, differed significantly in presence of the hetero species (GC+OC, GC+LT and GC+T) when compared to 8GC (Figure 3 and 4). GC+T had a significantly lower speed than 8GC (Dunnett’s test: *P* = 0.023; Figure 4b).

Interestingly, post-hoc tests revealed no significant differences in IID values (when compared to 8GC) across different treatments of starvation, space reduction and presence of hetero species (Figure 4a).

## 4. Discussion

In this study, we examined shoaling adaptations of GC with regards to different ecological and environmental manipulations. Our findings indicate that GC forms and maintains shoals by dynamically modulating the structural components, and these modulations vary across a range of tested manipulations. In absence of the manipulations, GC predominantly formed a single shoal (for both 4 and 8GC setups), however, the larger groups of 8GC had reduced speed and cohesion when compared to the smaller groups, the 4GC, possibly attributing to higher coordination costs in larger assemblages (Mallick et al. 2022). Further, to assess the shoaling adaptations under diverse conditions, we subjected the 8GC to different kinds of manipulations, ranging from presence of hetero species to starved conditions to reduction in habitat availability. Interestingly, in manipulative conditions, 8GC showed a large degree of modifications in different structural components, for example, a significant increase in shoal size and cohesive shoals, and significantly fewer shoal splits and lower number of solitary individuals, under both starved and space-reduced conditions, when compared to 8GC with no manipulations. Additionally, speed decreased significantly under space-reduced conditions and in presence of hetero species. Overall, the findings highlight the flexible adjustments of shoaling components in GC in response to different manipulative conditions, emphasizing the ability of the species to adapt to environmental challenges. This kind of behavioural plasticity possibly provides evolutionary advantages for the Cichlidae, a family that is largely known for their group living strategies (Taborsky, 2016; Jordan et al. 2021).

Studies have shown that animal sociality is highly dynamic in challenging situations, mostly reflecting the decision making and survival strategies of the species under severe conditions of stress (Hansen et al. 2015; Wilson et al. 2019; Kuntz et al. 2024). In our study, shoal size and cohesion in GC increased significantly (in comparison to control 8GC) under 24-hour starvation. Contrastingly, shoal splits and the number of solitary individuals decreased when compared to the control group (8GC), emphasizing an overall increase in coordination and sociality under the stress of starvation. Interestingly, similar shifts towards increased social cohesion have been observed in several species, referred to as a cooperative strategy adopted by species during challenging situations. For example, fire ants *Solenopsis invicta* Buren 1972 form raft-or ball-like structures during floods, increasing the group size and cohesion to enhance survival through collective buoyancy and protection (Foster et al. 2014). In the cooperatively breeding daffodil cichlid *Neolamprologus pulcher* (Trewavas and Poll 1952), juveniles from different parents formed more cohesive shoals in the absence of parental care compared to when parents were present (Zacke and Thünken, 2024), possibly strengthening their safety by increasing the numbers of nearest neighbours. Such cohesive large groups may also enhance the efficiency of information transfer in response to challenges (Ward et al. 2018). In our study, though an increase in cohesion was still maintained in GC_48H trials, the rest of the structural components, shoal size and splits, and number of solitary individuals, were not significantly different in GC_48H when compared to 8GC. Although in absolute terms, the trend in GC_48H was similar to that of GC_24H, both when compared to 8GC. These subtle differences in structural components of shoaling between GC_24H and GC_48H (both when compared to 8GC) can possibly be analogous to the strategies adopted by organisms when facing conditions of intermittent or acute stress (of starvation) versus long-term or chronic periods of stress (of starvation) (Kleinhappel et al. 2019). Nonetheless, under both the starvation regimes, GC consistently maintained high cohesion, demonstrating an increased tendency to cooperate to potentially contribute towards efficient transfer of information under challenging conditions (Ward et al. 2018).

The pattern of increased sociality was maintained even under the constrained condition of space reduction, wherein cohesion and shoal size increased, while shoal splits and solitary individuals decreased compared to the control group of 8GC. We speculate that this increased cohesion might not be an artefact of crowding (due to reduced space), as large area in the laboratory tank was still available for the shoaling fishes (ratio of area occupied by 8GC:total area available= 1:25.3) in GC_SR trials. However, unlike in starved conditions, GC also showed a reduction in speed under space-reduced conditions when compared to the control group of 8GC. Such reduced speed can also be associated with increased cohesion as documented by the shoals of *G. aculeatus*, who increased cohesion and decreased speed under conditions of high turbidity (MacGregor and Ioannou 2023). These findings under resource-constrained environments, for example under both starvation and space-reduction, suggest that GC are increasing their tendency to form social grouping, a strategy possibly adopted by the species to cope with challenges (Herbert-Read et al. 2017). Whether such an increased tendency towards sociality is a learned or an innate trait in GC, warrants future investigations while comparing their shoaling dynamics across different developmental stages (Magurran 1986; Buske and Gerlai, 2011).

Our study showed that GC participated in mixed-species shoals with T, LT, and OC. However, the mean percentage of splits increased, from approximately 19% in 8GC to nearly 48% and 63% in GC+T and GC+LT, respectively, and to about 42% in GC+OC, indicating a possible disruption or fission in shoals that included GC and either of the hetero species, the alien T or the closely related OC. Our results on documenting fission in mixed-species shoals are parallel with the findings by Desai and Ghosal (2024), where GC also showed a lower social preference towards both the hetero species, T and OC, when compared to significantly higher preference towards the conspecifics. Interestingly, in spite of the shoal splits, such mixed-species shoaling of native GC and alien *Oreochromis* spp. (T and LT) species could be a result of social interactions due to decades of co-occurrence in the same environment (Ward et al. 2003). Alternatively, this might be a strategy of *Oreochromis* spp., as shoaling with a native (familiar to the given environment) species (GC) may facilitate their invasion success (Camacho-Cervantes et al. 2014b) in a given ecosystem. GC+T also showed a significant reduction in speed when compared to 8GC. Studies have shown that mixed-species shoals are usually maintained via matching of swimming speeds among the shoaling species (Krause et al. 2005). For example, a study on Chinese bream *Parabramis pekinensis* Basilewsky 1855 and Qingbo *Spinibarbus sinensis* (Bleeker 1871) demonstrated that in mixed-species shoals, Chinese bream increased their swimming speed when compared to their speeds in the conspecific shoals (Tang et al. 2017). The authors speculated that increase in speed might have helped the bream to match the speed of *S. sinensis*, thereby maintaining shoal coordination. In our study, it seems like LT and OC have speeds comparable to that of GC, however, T might have a lower speed that possibly contributed towards an overall reduction in the shoaling speed of GC+T trials. Future studies should investigate the shoaling dynamics of *Oreochromis* spp. to characterize preferences and structural components (speed, shoal size, and NND) of *Oreochromis* spp. shoals, for better interpretation of the evolutionary advantages derived by the mixed-species shoals of GC and *Oreochromis* spp..

Our findings demonstrate that shoaling behaviour in GC is highly plastic, enabling rapid adjustments to varying environmental conditions. However, most of the adjustments took place in the local components (Allibhai et al. 2023), such as cohesion, speed, shoal size and occurrence of solitary fish. In contrast, the global parameter (Allibhai et al. 2023), IID, remained stable across all conditions, possibly highlighting its role in maintaining the overall stability of the shoal structures to facilitate collective coordination. Notably, GC displayed high levels of sociality under challenging conditions such as starvation and space reduction, suggesting the species (like many other cichlids; Zacke and Thünken, 2024) to be inherently social. Remarkably, even in the presence of *Oreochromis* spp., an alien and typically aggressive species (Jordan et al. 2020), GC formed mixed-species shoals. This behavioural flexibility points to the potential for such associations in natural habitats, where either the *Oreochromis* spp. may exploit the GC shoals for increased invasion success or GC may exhibit such adaptations in response to varying environmental pressures. Future studies should investigate the shoaling dynamics of *E. suratensis* in the wild to bridge the gap between manipulative laboratory conditions and complex, natural conditions. Such research would help clarify how ecological factors in the wild influence shoal formation, coordination, and species interactions, providing a deeper understanding of their adaptive strategies.

## Data availability

All the raw data and Python scripts used to analyse the data are available on Figshare: https://figshare.com/s/c20917098e4373b40070

## Author contributions

CD was responsible for conceptualization, investigation, data curation, formal analysis, and writing – original draft. RD contributed to formal analysis and validation. DN was responsible for python script development and formal analysis. RG contributed to funding acquisition, supervision, conceptualization, formal analysis, methodology, project administration, visualization, writing – original draft, and writing – review & editing.

## Acknowledgements

We are grateful to Suman Mallick for assistance with fish maintenance. The authors also acknowledge Bhavya Jariwala and Rashmita Ganguly for their help with video analysis. We acknowledge the University Grants Commission (UGC), Government of India, for providing fellowship support to CD.

